# Isogenic lines of rainbow trout (*Oncorhynchus mykiss)* as a tool to assess how growth and feeding behaviour are correlated to feed efficiency in fish

**DOI:** 10.1101/2023.01.10.523400

**Authors:** Simon Pouil, Thierry Kernéis, Edwige Quillet, Delphine Lallias, Laurent Labbé, Florence Phocas, Mathilde Dupont-Nivet

## Abstract

Accurately measuring individual feed intake is required to include feed efficiency (FE) as an objective in commercial breeding programs. Phenotyping individual feed intake through direct measurements remains complex in fish reared in groups. One way to overcome this challenge is to find proxies for estimating FE. This study aimed to investigate the correlations between fish FE and potential predictive criteria in rainbow trout *Oncorhynchus mykiss*. As predictive criteria, we considered the variations of body weight assessed as thermal growth coefficients (TGC) and the feeding behaviour assessed as the number of feed demands over a period and the proportion of the demands made before noon. Feed intake was assessed over three different periods in ten isogenic lines allowing a recording for each of the ten genotypes while keeping fish in groups. The protocol consisted of two successive feed deprivation and refeeding phases after initial basal growth periods. Correlations were then calculated between FE, measured either as feed conversion ratio (FCR) or residual feed intake (RFI), and the different indirect criteria. We found positive phenotypic correlations between FCR and RFI over the feed intake measurement periods (r ⩾ 0.79, *P* < 0.001). Assessment of the relationship between FE traits (FCR and RFI) calculated over the three measurement periods and TGC revealed no significant association. We found significant positive correlations between RFI calculated from the first feed intake measurement period and feeding behaviour, assessed as the number of feed demands (r = 0.42-0.49, *P* ⩽ 0.022). Nevertheless, such correlations were not established for the two other measurement periods following feed deprivation. While we demonstrated that the weight variations during feed deprivation are not a good proxy for FE (FCR or RFI) in rainbow trout, we also highlighted the interest in exploring more the relationships between FE and feeding behaviour in fish.

## 1. Introduction

Aquaculture is one of the fastest-growing animal protein supplies globally, while approximately 70% of aquaculture production requires external feed inputs (FAO, 2020). Nowadays, exogenous feeds for reared species are at the heart of concerns for the sustainability of finfish aquaculture. While exogenous feeds can represent up to 70% of farm costs (Doupé and Lymbery, 2004; de Verdal et al., 2018a), they are also responsible for environmental impacts directly resulting from feeds manufacturing and then, indirectly, by the release of nutrients by fish (Mente et al., 2006; Read and Fernandes, 2003). Furthermore, the mobilization of resources and arable lands for aquafeed production raises concerns about aquaculture’s social impact (Troell et al., 2014). Therefore, selecting fish turning feeds efficiently into meat is key to achieving the sustainable development of aquaculture.

In fish, feed efficiency (FE) has been traditionally measured by feed conversion ratio (FCR), defined as the ratio of feed intake (FI) to weight gain. However, FCR values can result from different biological conditions such as a reduced FI, an increase of biomass production or a combination of both (Cantalapiedra-Hijar et al., 2020). Thereby, FCR is mainly used for the zootechnical and economic performances assessment of a batch of animals at the farm level. An alternative to FCR is the residual feed intake (RFI), defined as the difference between the feed consumed by an animal and its expected consumption as predicted from a regression model involving the requirements for maintenance and growth through the integration of metabolic weight and weight gain over the recording period as the independent variables (Koch et al., 1963). Therefore, contrary to FCR (Besson et al., 2020; Kause et al., 2006; Thodesen et al., 1999), RFI is uncorrelated to body weight by construction and individuals with a negative RFI are more efficient than the mean of the population, while individuals with a positive RFI are less efficient.

Improvement of FE in fish can be achieved through various levers, including husbandry (El-Sayed, 2002; Yilmaz and Arabaci, 2010), nutrition (e.g., Xu et al., 2001; Bowyer et al., 2020), and genetic selection (e.g., de Verdal et al., 2018b, 2018a; Knap and Kause, 2018). Feed efficiency is now among the main traits of interest for fish breeding programs (Chavanne et al., 2016). However, except for family-based selection requiring heavy infrastructures and human labour, as previously done for Atlantic salmon in Norway (Gjedrem, 2010), selection for FE remains challenging, as the precise recording of individual FI is needed. Such phenotyping is challenging in fish. Indeed, fish are usually reared in large groups making it difficult to follow the FI of each individual. The usual way to measure FI in fish is to remove and count uneaten pellets allowing to calculate FI as the difference between the weight of feed given and the weight of feed uneaten by fish (wasted) (Jobling et al., 2001). Other approaches have been developed to measure individual FI in fish including the use of dyed feed, X-radiography and individual rearing (Besson et al., 2019; Jobling et al., 2001; Silverstein, 2006; Talbot and Higgins, 1983). An alternative method consisting of rearing small groups of fish in aquaria (∼5-15 fish together) with video recording has been proven to be efficient for FI determination in fish (de Verdal et al., 2019, 2018b, 2017; Just et al., 2021) while developments are currently performed to automate video analysis by artificial intelligence (Zhou et al., 2018). Recently, Dvergedal et al. (2022) suggested that lipid deposition in rainbow trout *Oncorhynchus mykiss*, which is related to FE, could be phenotyped through the C stable isotope contents in feeds and fish. Nevertheless, all of these methods have limitations, including measurement inaccuracies with a lack of repeatability and/or measurement conditions too far from production conditions (reviewed by de Verdal et al. (2018a)) while the most recent ones are still requiring methodological developments.

Because collecting individual FI data in fish through direct measurements remains complex, many indirect criteria such as traits derived: 1) from bio-energetic models: lipid deposition, nitrogen retention, oxygen consumption and 2) from growth models: growth and changes in the body after starvation and refeeding periods, have been studied in recent years for estimating FE. Focusing on body weight loss during starvation and gain during the subsequent refeeding period, Grima et al. (2010a, 2010b) suggested that selection for European seabass *Dicentrarchus labrax* losing less weight during the starvation period should improve FE and increase muscle fatness. However, in rainbow trout, Grima et al. (2008) did not find any significant correlation between RFI and weight variations during starvation or refeeding periods. However, they highlighted that an index combining growth performances over all the experimental periods explained up to 59% of RFI variations.

Feed efficiency can also be linked to feeding behaviour. Indeed, several authors have shown that in terrestrial-farmed animals, feed-efficient individuals are less active while feeding (e.g., Kelly et al., 2010; Montanholi et al., 2010). In fish, Martins et al. (2006) have shown that African catfish *Clarias gariepinus* with lower RFI spend less time eating. In Nile tilapia *Oreochromis niloticus*, Martins et al. (2011a) found that RFI was negatively correlated with feeding latency (i.e. the time each fish takes to consume the first pellet), while positive correlations were found with the total feeding time and the number of feeding acts. Nevertheless, such relationships have been poorly investigated in the literature and, based on the limited information available, seem to be contrasted depending on the size of the fish used (Martins et al., 2011b).

On an experimental scale, one tool to overcome the drawbacks of the usual FI assessment methods in fish is using isogenic lines (Komen and Thorgaard, 2007a; Quillet et al., 2007b; Franěk et al., 2020). In such lines, all individuals are genetically identical. Crossing individuals from two homozygous isogenic lines allows to obtain genetically identical but heterozygous individuals in the next generation (de Verdal et al., 2018a). Although isogenic lines are produced for experimental purposes only, they are an excellent tool for studying the variability of FE, which is highly sensitive to environmental variations. Because the genetic variability within an isogenic line is null, fish belonging to the same line reared in the same tank allow a precise assessment of FI of an individual genotype while maintaining social interactions between individuals for each of the genotypes. In addition, combining information from different isogenic lines gives access to the genetic variability of FE.

The aims of this study were: (1) to confirm the existence of genetic variability in FE, (2) to estimate correlations between RFI, FCR and body weight variations during successive FD and CG periods and (3) to investigate the link between feeding behaviour and FE in rainbow trout.

## 2. Materials and Methods

### 2.1. Ethical statement

All the experiments were carried out at the INRAE experimental facilities (PEIMA, INRAE, 2021, Fish Farming systems Experimental Facility, DOI: 10.15454/1.5572329612068406E12, Sizun, France) authorised for animal experimentation under the French regulation D29-277-02. The experiments were carried out from January 2008 to July 2009 following the Guidelines of the National Legislation on Animal Care of the French Ministry of Research (Decree N°2001-464, May 29, 2001). At this date, project authorisation was not required. Experiments were conducted under the official license of M. Dupont-Nivet (A29102).

### 2.2. Heterozygous isogenic lines

Experimental fish were produced and reared in the INRAE experimental fish farm (PEIMA, Sizun, France) as described by Millot et al. (2014) and Lallias et al. (2017). Parents were issued from the INRAE homozygous isogenic lines. These lines were previously established after two generations of gynogenesis and further maintained within each line by single-pair mating using sex-reversed XX males (Quillet et al., 2007b). Mating 24 homozygous females produced ten heterozygous isogenic lines from a single maternal isogenic line (spawning on the same day with similar egg weights) with ten sex-reversed XX males from ten other homozygous isogenic lines. Eggs were mixed and then divided into ten batches, each batch being fertilised by one of the ten homozygous males. To avoid confusion with the INRAE homozygous isogenic lines, heterozygous lines will be named the line’s original name where the sire came from, plus “h” as heterozygous.

Eggs were mixed and then randomly divided into ten batches, each fertilised by milt from one of the ten sires. Since there is only one maternal line, progeny differs only through paternal genetic effects across batches. Fertilised eggs were incubated at 11.4°C. At the eyed stage, 1500 eggs of each of the ten produced heterozygous lines were distributed in 0.25 m^3^ indoor tanks (n = 3 per line, 500 fish per replicate) supplied with natural spring water at the same temperature. Fish were transferred in 1.8 m^3^ outdoor tanks (n=3 per line) supplied with dam water (11.3-16.0°C) after 137 days post-fertilization (dpf) up to the end of the experiment (548 dpf). In order to keep a density below 50 kg m^-3^, regular random eliminations were performed, and 100 fish per replicate remained at 548 dpf. Commercial pellets (BioMar’s Biostart range and Le Gouessant’s B-Mega range) were distributed by automatic feeders during the indoor phase and self-feeders (Imetronic®, France) during the outdoor rearing phase (see Section 2.4.2 for details). Feed ration was maintained *ad libitum* throughout the experiment. Mortality was checked every day.

### 2.3. Successive experimental phases

The experiment reported here started at the beginning of the outdoor rearing phase (*i*.*e*. 137 dpf). The experimental protocol is summarised in Figure 1. After two basal growth periods separated by chronic stress (see Lallias et al. (2017) for details), the protocol consisted of two repeated phases of FD and refeeding periods. Following an FD period, fish tend to compensate for the loss of growth experienced during the FD period by increasing their growth more than usual, a phase known as the CG period (Ali et al., 2003).

**Figure 1.**
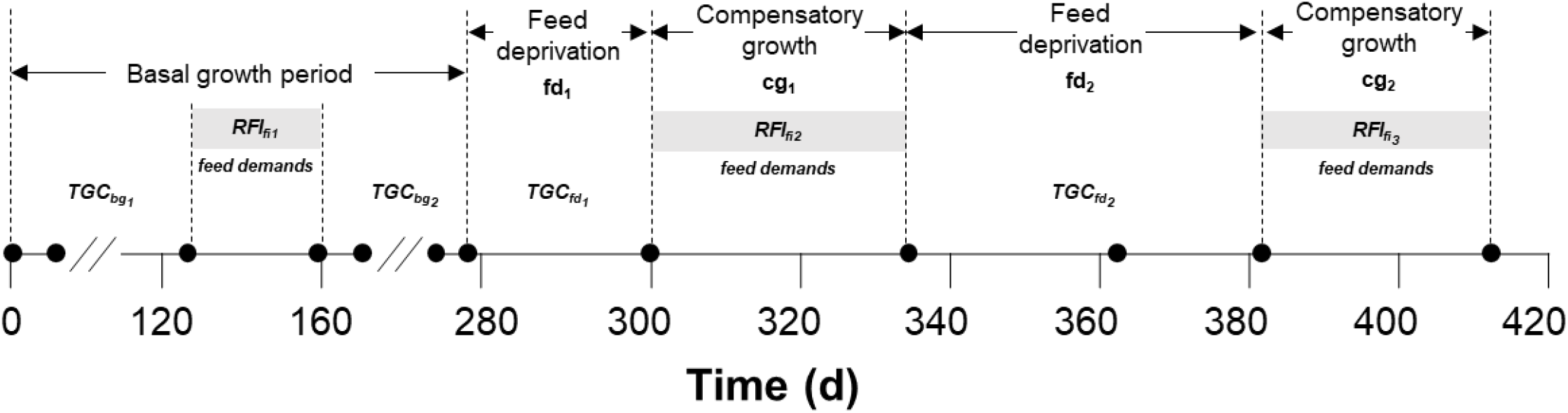
Schematic outline and time frame of the experiment performed using ten rainbow trout isogenic lines. All the different experimental periods are indicated: basal growth periods (bg_1_ and bg_2_), feed deprivations periods (fd_1_ and fd_2_) and compensatory growth periods (cg_1_ and cg_2_) and the different traits recorded as well: Thermal Growth Coefficients (TGC), Residual Feed Intake (RFI) and feed demands.

The experiment was divided into six different phases: two basal growth periods (bg_1_; from day 1 to day 129, *i*.*e*. 128 d and bg_2_; from day 186 to day 277, *i*.*e*. 91 d), FD period 1 (fd_1_; from day 278 to day 300, *i*.*e*. 22 d), CG period 1 (cg_1_; from day 301 to day 335, *i*.*e*. 34 d), FD period 2 (fd_2_; from day 336 to day 387, *i*.*e*. 48 d), and CG period 2 (cg_2_; from day 388 to day 415, *i*.*e*. 29 d). Feed intake (FI) was measured three times during the experiment, as shown in Figure 1. The first period of FI assessment was carried out from day 129 to day 161 (*i*.*e*. 32 d) to provide FE of the tested isogenic lines under normal growth conditions. The period from day 162 to day 185 was not considered here because fish were exposed to stress challenges which strongly influenced their growth (Figure 2). Then, the other FI measurement periods (fi) were performed during cg_1_ and cg_2_.

**Figure 2.**
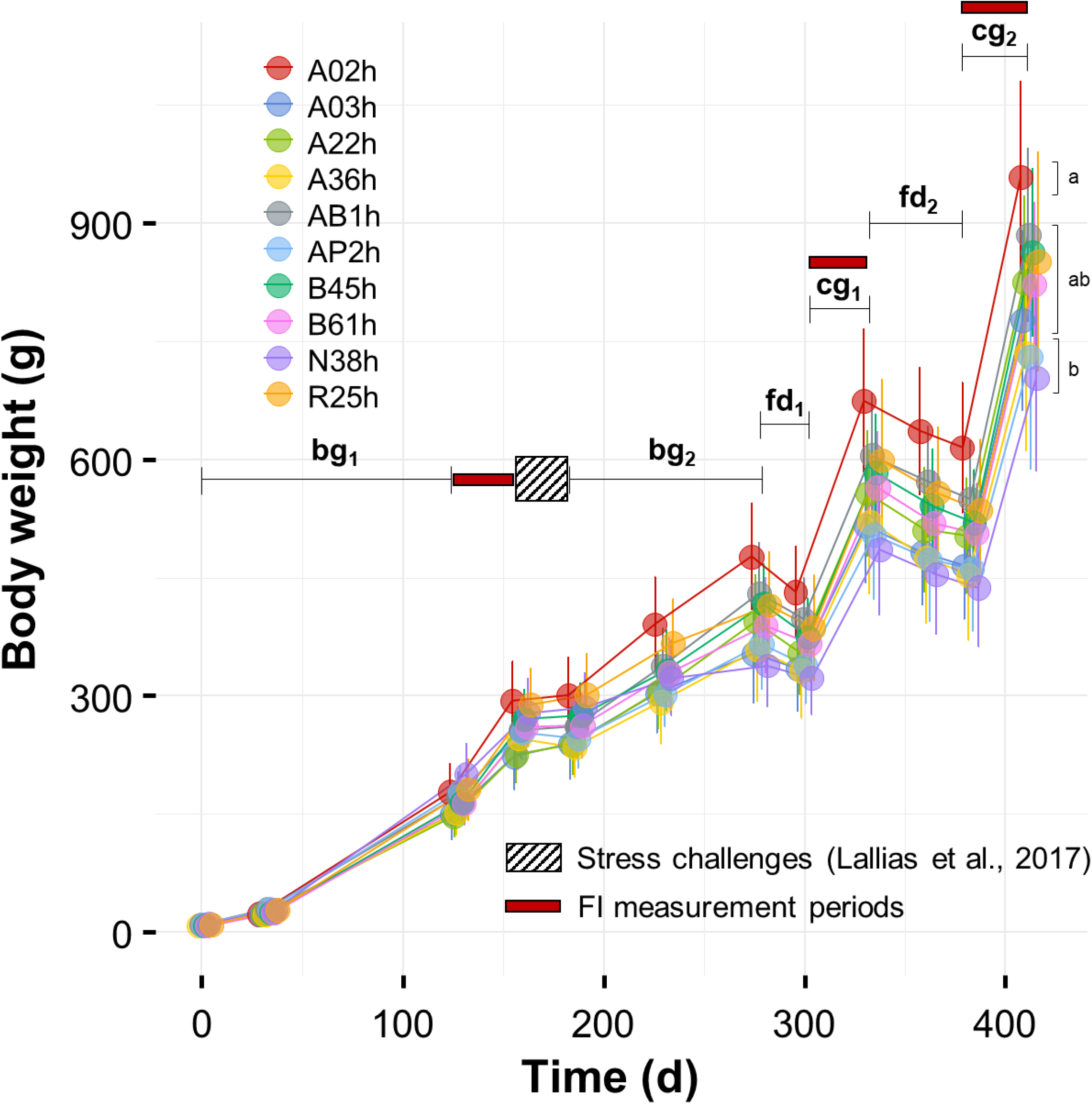
Growth kinetics of the ten isogenic lines over the experiment. Values are Means ± SD (n = 3). The different experimental periods are indicated in black: basal growth periods (bg_1_ and bg_2_), FD periods (fd_1_ and fd_2_) and CG periods (cg_1_ and cg_2_). Letters denote significant differences in final weights between lines.

### 2.4. Traits measured

#### 2.4.1. Body weight variations

Body weight was measured at the beginning and then 11 times throughout the experiment (n ≈ 100 fish per replicate). Fish were starved for 24h before any sampling. Weight gain or loss in each period of interest (*i*.*e*. bg_1_, bg_2_, fd_1_, cg_1_, fd_2_, cg_2_) was expressed as thermal growth coefficient (TGC), offering a standardised measure of growth that is unaffected by body weight, time interval, and water temperature (Iwama and Tautz, 1981; Grima et al., 2010b). TGC was calculated according to the following equation:

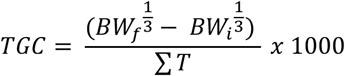

where BW_f_ and BW_i_ are the final and initial average body weights of the considered period, and ΣT is the sum of daily temperatures. Growth rates are referred to as TGC_bg1_, TGC_bg2,_ TGC_fd1,_ TGC_cg1,_ TGC_fd2_ and TGC_cg2_ for the different separated periods.

#### 2.4.2. Feed efficiency and feed demands

Self-feeders were used throughout the experiment to characterise the feeding behaviour of each isogenic line. This feeding technique is based on the learning ability of fish, and feed is delivered depending on their demand (Azzaydi et al., 2007). In a self-feeding system, fish are assumed to precisely control the feed distribution by activating a “trigger” sensor (Mambrini et al., 2004; da Silva et al., 2016). In order to limit involuntary demands and subsequent waste, sensors were placed just above the water. In addition, the self-feeders were configured to make access to the pellets more difficult according to the quantity of feed distributed over the day by increasing the inhibition time of the probe and by requiring two triggerings for pellet distribution. The protocol was adjusted over time according to the number of demands and the fish appetite. All the self-feeders were connected to a computer system (Imetronic®, France), allowing a continuous recording of all the fish’s feed distribution (sorted as rewarded demands when they led to feeding or unrewarded demands if not) during the FI measurement periods.

Two indicators of FE were used. The first one was the feed conversion ratio calculated as follows:

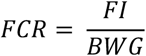

where FI is the feed intake and BWG is the body weight gain. Despite the simplicity of calculation and interpretation, one of the problems with FCR lies in the fact that the ratio obtained can result from different biological conditions: (1) a decrease in ingestion, (2) an increase in growth or (3) a combination of the two (Cantalapiedra-Hijar et al., 2020). Other criteria have been proposed in animal genetic selection to overcome the problems associated with ratios. The most frequently used criterion by geneticists is the residual feed intake (RFI; Koch et al., 1963), which was used as the second indicator for FE. RFI is calculated according to the following equation:

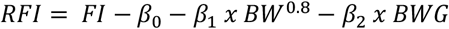

with β_0_ the regression intercept and β_1_ the partial regression coefficient of the animal’s FI on metabolic body weight. Average weights over the measurement period 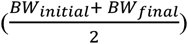 are scaled to metabolic weights (BW^0.8^; Clarke and Johnston, 1999; Jobling, 2002), while β_2_ is the partial regression coefficient of the animal’s FI on BWG. This model allocates the FI according to expected maintenance and growth requirements, the remaining part being defined as the RFI (Koch et al., 1963; de Verdal et al., 2018a).

### 2.5. Statistical analyses

The significance level for statistical analyses was set to α = 0.05. All statistics were performed using R freeware version 4.3.1 (R Development Core Team, 2020). The final body weight was compared between lines using a linear mixed model with “line” as a fixed effect and “replicate” as a random effect. The model was fitted using the *nlme* package, and contrasts were analysed using the *emmeans* package. Assumptions of normality and homoscedasticity were checked by visual inspections of residuals against fitted values.

Differences in weight variations, TGC, FI and RFI estimated for each experimental period between lines were tested using the aligned rank transformation for nonparametric factorial analysis (*art*(), *ARTool* package) using the tanks as the experimental units. Contrasts were computed using the function *art*.*con*() from the same R package.

Regarding the feeding behaviour data, for each of the three FI measurement periods, the number of daily feed demands was compared between lines as described above for final weight. The daily feeding profile (% demand activity h^-1^ per day) for each FI measurement period was compared between lines with a generalised linear mixed model with “line” and “hour” as fixed effects while “replicate” was included as a random effect. The model was fitted using the *glmmTMB* package allowing to fit generalised linear mixed models with various extensions.

The correlations between the variables (averaged by tank for rewarded and unrewarded feed demands) were examined with the *rcorr*() function of the *Hmisc* package.

## 3. Results and Discussion

### 3.1. Effects of feed deprivation on survival and growth

As expected, fish overcame starvation, and the two FD periods did not affect survival in mortality rate (mortality rate < 1% over fd_1_ and fd_2_). Weatherley and Gill (1981) already highlighted the ability of rainbow trout to deal with prolonged starvation (*i*.*e*. up to 13 weeks) without deleterious effect on subsequent zootechnical performances. The final individual weights by line after the 415-d experiment (*i*.*e*. 552 dpf) ranged from 704 ± 119 g to 958 ± 123 g for N38h and A02h, respectively (Figure 2), while TGC significantly differed between the lines over the experiment (*P* < 0.001). The ten lines differed slightly for response to FD, with an average weight loss ranging from 5 to 10% during the first FD period (fd_1_) vs from 8 to 13% during the second FD period (fd_2_), with significant differences recorded among the lines for both periods (*P* <0.002). These results confirm the existence of genetic variability in response to feed deprivation (Grima et al., 2008) and highlight differences in the requirements for maintenance between lines. Thermal growth coefficients estimated from the two FD periods were always highly correlated with weight loss (r ⩾ 0.96, *P* < 0.001; Table 1) and were significantly different between lines (TGC_fd1_: *P* < 0.001, and TGC_fd2_: *P* = 0.001; Table 2). The correlation between TGC recorded during the two FD periods was low (TGC_fd1_ and TGC_fd2_; r = 0.19, *P =* 0.326; Table 1 and Figure 3). For the first CG period, the TGC values ranged from 0.60 to 0.75 with no significant differences between lines (TGC_cg1_, *P* = 0.121; Table 2), while TGC_cg2_ values were 2 to 3 times lower with significant differences between lines (*P* = 0.007; Table 2). In contrast, TGC recorded during the two CG periods was moderately but significantly correlated (TGC_cg1_ and TGC_cg2_; r = 0.39, *P =* 0.035; Table 1). Overall, the lines that lost the least weight during the FD periods are those that gained the most during the CG periods, with a significant negative correlation observed between TGC_fd1_ and TGC_cg1_ (r = -0.53, *P =* 0.003; Table 1). These findings are in accordance with Dupont-Prinet et al. (2010) showing that there was a negative correlation between body weight losses during FD and body weight gain during CG in European sea bass.

**Table 1.** Correlations between body weight variations (BWG; %) over FD periods (fd_1_ and fd_2_) and CG periods (cg_1_ and cg_2_) and Thermal Growth Coefficients (TGC) calculated for the same experimental periods. Significant correlations are highlighted in bold.

|  | BWG<sub>fd1</sub> | TGC<sub>fd1</sub> | BWG<sub>cg1</sub> | TGC<sub>cg1</sub> | BWG<sub>fd2</sub> | TGC<sub>fd2</sub> | BWG<sub>cg2</sub> | TGC<sub>cg2</sub> |
| --- | --- | --- | --- | --- | --- | --- | --- | --- |
| BWG<sub>fd1</sub> |  | - | - | - | - | - | - | - |
| TGC<sub>fd1</sub> | **0.99** |  | - | - | - | - | - | - |
| BWG<sub>cg1</sub> | **-0.38** | **-0.37** |  | - | - | - | - | - |
| TGC<sub>cg1</sub> | **-0.49** | **-0.53** | **0.90** |  | - | - | - | - |
| BWG<sub>fd2</sub> | 0.09 | 0.05 | -0.15 | -0.02 |  | - | - | - |
| TGC<sub>fd2</sub> | 0.20 | 0.19 | -0.23 | -0.21 | **0.96** |  | - | - |
| BWG<sub>cg2</sub> | -0.02 | 0.04 | 0.16 | -0.04 | -0.23 | -0.14 |  | - |
| TGC<sub>cg2</sub> | -0.27 | -0.28 | 0.34 | **0.39** | -0.04 | -0.12 | **0.80** |  |

**Table 2.** Thermal growth coefficient (TGC) in 10 rainbow trout isogenic lines during two basal growth periods (bg_1_ and bg_2_), followed by two periods of FD (fd_1_ and fd_2_) alternated with two periods of CG (cg_1_ and cg_2_) and RFI calculated from the three feed intake measurement periods (fi_1_, fi_2_ and fi_3_). Values are Means ± SD. Letters denote significant differences (p < 0.05).

| Traits | Isogenic lines |  |  |  |  |  |  |  |  |  |
| --- | --- | --- | --- | --- | --- | --- | --- | --- | --- | --- |
|  | A02h | A03h | A22h | A36h | AB1h | AP2h | B45h | B61h | N38h | R25h |
| Thermal growth coefficients (TGC) |  |  |  |  |  |  |  |  |  |  |
| bg<sub>1</sub> | 2.00 ± 0.10<sup>a</sup> | 1.81 ± 0.16<sup>a</sup> | 1.80 ± 0.04<sup>a</sup> | 1.80 ± 0.03<sup>a</sup> | 1.87 ± 0.11<sup>a</sup> | 1.94 ± 0.15<sup>a</sup> | 1.88 ± 0.06<sup>a</sup> | 1.89 ± 0.11<sup>a</sup> | 1.99 ± 0.11<sup>a</sup> | 1.91 ± 0.08<sup>a</sup> |
| bg<sub>2</sub> | 1.95 ± 0.07<sup>a</sup> | 1.54 ± 0.10<sup>bc</sup> | 2.01 ± 0.35<sup>ab</sup> | 1.65 ± 0.26<sup>abc</sup> | 2.04 ± 0.15<sup>a</sup> | 1.58 ± 0.23<sup>abc</sup> | 1.71 ± 0.12<sup>abc</sup> | 1.61 ± 0.10<sup>abc</sup> | 0.67 ± 0.22<sup>c</sup> | 1.33 ± 0.28<sup>c</sup> |
| fd<sub>1</sub> | -1.49 ± 0.07<sup>ab</sup> | -0.87 ± 0.09<sup>c</sup> | -1.58 ± 0.38<sup>ab</sup> | -1.15 ± 0.35<sup>abc</sup> | -1.17 ± 0.16<sup>abc</sup> | -1.15 ± 0.26<sup>abc</sup> | -1.62 ± 0.22<sup>a</sup> | -0.97 ± 0.26<sup>bc</sup> | -0.72 ± 0.23<sup>c</sup> | -1.09 ± 0.10<sup>abc</sup> |
| cg<sub>1</sub> | 3.94 ± 0.22<sup>a</sup> | 3.52 ± 0.29<sup>a</sup> | 3.73 ± 0.20<sup>a</sup> | 3.66 ± 0.06<sup>a</sup> | 3.58 ± 0.03<sup>a</sup> | 3.18 ± 0.48<sup>a</sup> | 3.73 ± 0.31<sup>a</sup> | 3.61 ± 0.11<sup>a</sup> | 3.25 ± 0.52<sup>a</sup> | 3.73 ± 0.17<sup>a</sup> |
| fd<sub>2</sub> | -0.49 ± 0.03<sup>ab</sup> | -0.51 ± 0.02<sup>ab</sup> | -0.51 ± 0.01<sup>ab</sup> | -0.68 ± 0.15<sup>a</sup> | -0.50 ± 0.08<sup>ab</sup> | -0.44 ± 0.03<sup>b</sup> | -0.59 ± 0.02<sup>a</sup> | -0.55 ± 0.05<sup>ab</sup> | -0.50 ± 0.01<sup>ab</sup> | -0.58 ± 0.05<sup>a</sup> |
| cg<sub>2</sub> | 3.52 ± 0.07<sup>ab</sup> | 3.75 ± 0.15<sup>a</sup> | 3.72 ± 0.17<sup>ab</sup> | 3.50 ± 0.09<sup>ab</sup> | 3.66 ± 0.10<sup>ab</sup> | 3.30 ± 0.49<sup>ab</sup> | 3.86 ± 0.23<sup>a</sup> | 3.64 ± 0.09<sup>ab</sup> | 3.40 ± 0.09<sup>b</sup> | 3.53 ± 0.07<sup>ab</sup> |
| Feed intake (g kg<sup>-1</sup> d<sup>-1</sup>) |  |  |  |  |  |  |  |  |  |  |
| fi<sub>1</sub> | 14.4 ± 0.81<sup>ab</sup> | 13.5 ± 2.39<sup>ab</sup> | 12.1 ± 2.11<sup>ab</sup> | 15.5 ± 1.15<sup>ab</sup> | 11.7 ± 2.78<sup>ab</sup> | 11.8 ± 1.94<sup>ab</sup> | 16.3 ± 1.73<sup>a</sup> | 15.0 ± 0.50<sup>ab</sup> | 10.2 ± 1.63<sup>b</sup> | 14.0 ± 0.84<sup>ab</sup> |
| fi<sub>2</sub> | 14.6 ± 0.96<sup>a</sup> | 14.9 ± 1.74<sup>a</sup> | 13.6 ± 0.64<sup>a</sup> | 16.6 ± 0.82<sup>a</sup> | 13.3 ± 0.54<sup>a</sup> | 12.3 ± 1.94<sup>a</sup> | 14.1 ± 1.41<sup>a</sup> | 13.5 ± 0.31<sup>a</sup> | 13.4 ± 3.12<sup>a</sup> | 15.8 ± 0.95<sup>a</sup> |
| fi<sub>3</sub> | 20.5 ± 0.80<sup>a</sup> | 24.6 ± 1.55<sup>a</sup> | 22.7 ± 2.07<sup>a</sup> | 23.9 ± 1.66<sup>a</sup> | 21.8 ± 0.95<sup>a</sup> | 20.7 ± 4.37<sup>a</sup> | 23.6 ± 1.81<sup>a</sup> | 22.6 ± 1.75<sup>a</sup> | 22.7 ± 0.71<sup>a</sup> | 22.4 ± 0.90<sup>a</sup> |
| Feed conversion ratio (FCR) |  |  |  |  |  |  |  |  |  |  |
| fi<sub>1</sub> | 0.78 ± 0.05<sup>a</sup> | 0.94 ± 0.06<sup>a</sup> | 0.79 ± 0.01<sup>a</sup> | 0.83 ± 0.06<sup>a</sup> | 0.82 ± 0.08<sup>a</sup> | 0.90 ± 0.05<sup>a</sup> | 0.83 ± 0.08<sup>a</sup> | 0.82 ± 0.05<sup>a</sup> | 0.88 ± 0.08<sup>a</sup> | 0.81 ± 0.02<sup>a</sup> |
| fi<sub>2</sub> | 0.86 ± 0.02<sup>abc</sup> | 0.90 ± 0.02<sup>ab</sup> | 0.79 ± 0.02<sup>c</sup> | 0.96 ± 0.10<sup>ab</sup> | 0.84 ± 0.03<sup>abc</sup> | 0.84 ± 0.03<sup>abc</sup> | 0.83 ± 0.02<sup>abc</sup> | 0.82 ± 0.03<sup>ac</sup> | 0.87 ± 0.07<sup>abc</sup> | 0.94 ± 0.03<sup>b</sup> |
| fi<sub>3</sub> | 0.95 ± 0.02<sup>a</sup> | 0.95 ± 0.03<sup>a</sup> | 0.91 ± 0.02<sup>a</sup> | 0.99 ± 0.02<sup>a</sup> | 0.93 ± 0.03<sup>a</sup> | 0.92 ± 0.10<sup>a</sup> | 0.93 ± 0.04<sup>a</sup> | 0.94 ± 0.05<sup>a</sup> | 0.96 ± 0.04<sup>a</sup> | 0.99 ± 0.01<sup>a</sup> |
| Residual feed intake (RFI) |  |  |  |  |  |  |  |  |  |  |
| fi<sub>1</sub> | -2.85 ± 4.02<sup>a</sup> | 4.18 ± 0.27<sup>a</sup> | -4.92 ± 2.50<sup>a</sup> | 1.53 ± 5.02<sup>a</sup> | -2.97 ± 5.08<sup>a</sup> | 2.09 ± 3.45<sup>a</sup> | 3.47 ± 8.08<sup>a</sup> | 0.78 ± 4.36<sup>a</sup> | -0.58 ± 3.83<sup>a</sup> | -0.72 ± 1.80<sup>a</sup> |
| fi<sub>2</sub> | -3.41 ± 4.74<sup>abcd</sup> | 5.73 ± 3.76<sup>abc</sup> | -15.38 ± 4.15<sup>d</sup> | 18.58 ± 19.00<sup>ac</sup> | -4.74 ± 5.81<sup>abcd</sup> | -2.57 ± 4.03<sup>abcd</sup> | -7.05 ± 3.34<sup>abd</sup> | -9.49 ± 5.82<sup>bd</sup> | 1.78 ± 11.12<sup>abcd</sup> | 16.56 ± 8.07<sup>c</sup> |
| fi<sub>3</sub> | -0.90 ± 5.30<sup>a</sup> | 2.49 ± 8.45<sup>a</sup> | -10.39 ± 5.96<sup>a</sup> | 13.29 ± 5.65<sup>a</sup> | -6.57 ± 9.88<sup>a</sup> | -3.17 ± 21.89<sup>a</sup> | -6.91 ± 11.63<sup>a</sup> | -3.26 ± 15.71<sup>a</sup> | 4.70 ± 9.75<sup>a</sup> | 10.73 ± 3.03<sup>a</sup> |

**Figure 3.**
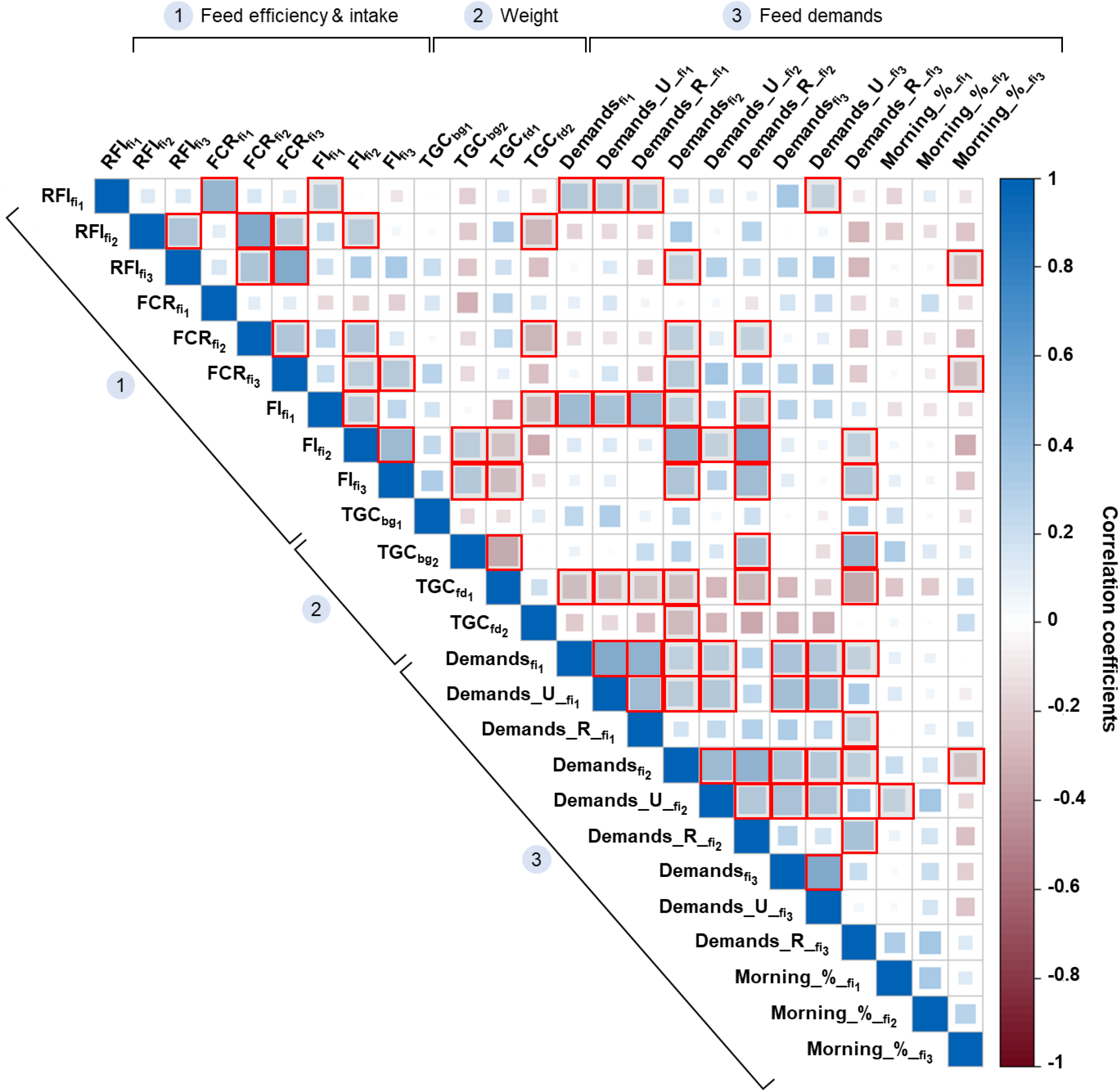
Correlation coefficients of all the measured phenotypes related to (1) Feed efficiency (FCR and RFI) and feed intake (FI) estimated during the three feed intake measurements periods (fi); (2) Weight assessed as TGC over all the experimental periods (bg, fd and cg); and 3) Feed demands over the three feed intake measurements periods considering all the demands (“Demands”), the unrewarded demands (“Demands_U”), the rewarded demands (“Demands_R”) or the proportion of the demands made before noon (“Morning_%”). The size and the colour gradient of the squares denote the intensity of the correlations. Significant correlations (*p* < 0.05) are highlighted in red.

The differing level of response between the periods may also be the consequence of rapid fish adaptation to a repeated cycle of feed deprivation and refeeding (Blake et al., 2006), with modulation of their physiological responses as suggested by Grima et al. (2008).

### 3.2. Feed intake and feed efficiency measurements

From a practical point of view, for each isogenic line, keeping fish in a group allowed us to get values of FI (see Table 2) from at least 29 consecutive days. This period exceeds the number of meals needed to obtain a reliable mean value of FI in fish; de Verdal et al., 2018b) while maintaining social interactions between individuals for each genotype. In this study, measurements of FI were performed either during the basal growth period (fi_1_) or in CG periods (fi_2_ and fi_3_). We observed that the increase in FI between fi1 and fi2 was, overall, low and line dependent, while FI_fi3_ was considerably higher in the ten lines (Table 1). Most studies agree that compensatory growth in fishes is enabled by an increase in FI, even if the pattern of FI differs between species (Wu et al., 2002). The effective improvement of FE during compensatory growth is debatable. It was demonstrated in rainbow trout (Nikki et al., 2004) and pikeperch *Sander lucioperca* (Mattila et al., 2009) that compensatory growth was permitted by increasing FI but not an improvement of FE. In contrast, Mambrini et al. (2004) in rainbow trout and Oh et al. (2007) in red sea bream *Pagrus major* observed both an improvement of FE measured as the inverse of FCR and a rise of FI during refeeding in fasted fish compared to control unfasted fish.

We found significant differences in FCR and RFI between lines only in fi_2,_ while variability in FCR and RFI remains high among the replicates of a given line (Table 2). Over the second FI measurement period (*i*.*e*. fi_2_), the line A22h was the most feed-efficient with values of FCR and RFI of 0.79 ± 0.02 and -15.38 ± 4.15, respectively, while A36h was the less feed-efficient line with FCR of 0.96 ± 0.10 and RFI of 18.58 ± 19.00 (Table 2). Few studies investigated the correlations between FCR and RFI in fish. Overall, we found high phenotypic correlations between FCR and RFI measured over the same period, with a slightly lower correlation during the first period (fi_1_; r = 0.79, *P* < 0.001; Figure 3) and very high during the last two measurement periods (fi_2_ and fi_3_; r = 0.99 and 0.97, *P* < 0.001; Figure 3). These results are in line with the literature on rainbow trout. Indeed, Kause et al. (2016) found strong positive phenotypic correlations between FCR and RFI in fish older than two years. Based on the data from Grima (2010) and Grima et al. (2008), we were able to recalculate a positive phenotypic correlation between FCR and RFI in isogenic lines of rainbow trout with fish of comparable size to the individuals used in our experiment. Using ten different isogenic lines, we confirmed genetic-based variation of FE in rainbow trout, a requirement to implement breeding programs. However, such variations are not consistent over time, presumably due to the feeding practice. Indeed, while previous works argued that self-feeding minimises competition for feed because feed is more accessible (Boujard et al., 2002; Mambrini et al., 2004), the high variability in FCR and especially RFI between replicates of a given isogenic line we observed suggested that social interactions can strongly affect FE even when fish are fed using self-feeders. Indeed, within the same rearing unit, FI is not necessarily homogeneous among fish, and this can be exacerbated using self-feeders where a few high-triggering fish can drive group feeding behaviour, as shown in European sea bass and such social interactions can change over time (Millot and Bégout, 2009).

Overall, the correlations between the three FCR and the three RFI values derived from the three FI periods were, as shown in Figure 3, low between FCR _fi1,_ RFI_fi1_ and the two other measurements (FCR_fi2_ and FCR_fi3_; r = 0.11 and 0.12, *P* ⩾ 0.450, RFI_fi2_ and RFI_fi3_; r = 0.14, *P* ⩾ 0.468). Interestingly, the correlations were significant only between the last two FCR and the last two RFI measured during CG periods (*r* = 0.53 and 0.55, *P* ⩽ 0.003). Such inconsistency over time can be explained by: (1) the difference in age of the fish and/or (2) the fact that the two last FI measurements periods occurred after feed deprivation in large fish (⩾ 350 g) while the first FI period was done during a basal growth period in smaller fish (⩽ 200 g). Here, the phenotypic correlations established between the FCR and RFI measured at three different periods indicate that these traits are reliable for phenotyping FE but may require repeated measurements. These results suggest that our measurement methodology, allowing FI recording over several consecutive weeks, exhibits relatively high repeatability. Nevertheless, although of high experimental interest, the acquisition of individual FI value for each genotype from fish kept in groups is conditioned by the use of isogenic lines and, therefore, this method is not directly applicable in a breeding program.

### 3.3. Correlations between feed efficiency and weight variations

We did not find significant phenotypic correlations between TGC recorded in basal growth periods (TGC_bg1_ and TGC_bg2_) and FE traits (FCR and RFI). In livestock species, FE is generally correlated with growth, but this is still debatable in fish. In rainbow trout, while Silverstein (2006) estimated negative phenotypic correlations between TGC and RFI (r = -0.57 to -0.31), Kause et al. (2016) found no phenotypic correlations between daily weight gain (DWG) and RFI (r = 0.08). Interestingly, strong genetic correlations have been highlighted between DWG and FCR (r = -0.63 ± 0.30; Kause et al., 2016) and BW and inverse of FCR (r = 0.63-0.99; Henryon et al., 2002). Nevertheless, no significant genetic correlations have been found between DWG and RFI (Kause et al., 2016).

In practice, fish breeders expect to improve FE by selecting the fasted growing fish, hypothesising that faster-growing fish will be more efficient (Besson et al., 2020; Kause et al., 2016; Knap and Kause, 2018). Still, few selection response studies have investigated the correlated response in FE to selection for growth and provided contrasting results. Recently, Vandeputte et al. (2022) estimated realised genetic gains on FCR and other traits in a commercial population of rainbow trout selected for improved growth, carcass yield and fillet fat over ten generations. The authors demonstrated significant improvement in FCR even if the trait was not directly included in the breeding goal.

We based our study on previous works showing that performance traits (i.e. weight variations) during successive FD periods can be correlated to FE in fish (e.g. Grima et al., 2010; Li et al., 2005). In our experiment, an assessment of the relationship between FE traits (FCR and RFI) calculated over the three FI measurement periods and growth-related traits (TGC) revealed no significant correlation. The only exception was the significant negative correlations between TGC_fd2_ and FCR_fi2_ or RFI_fi2_ with respective r = -0.49 and -0.47 (*P* ⩽ 0.006). These results suggest that the fish that had lost the least weight during the first phase of FD were the most feed efficient during the following CG period (fi2). Using weight loss during FD periods as an indirect criterion to select FE required that it is genetically correlated to RFI, the most interesting FE trait in selection (see Section 2.4.2). Here, we found limited phenotypic correlations between the TGC measured during FD periods and RFI, raising questions about the interest of such a criterion to evaluate FE in rainbow trout. Indeed, based on our estimates of phenotypic correlations, we can reasonably assume that the genetic correlations between these traits are probably weak. Nevertheless, in Nile tilapia, although de Verdal et al. (2018b) found a weak phenotypic correlation between RFI and TGC during the FD period (r = 0.09), the genetic correlation was high (r = 0.70), indicating that TGC during FD period could be a promising selection criterion for FE in this species. Such a relationship needs to be confirmed in rainbow trout, and based on current knowledge, there is no strong evidence of the interest in using such an indirect selection criterion for FE in rainbow trout.

### 3.4. Feeding behaviour and its link with FE

One of the originalities of this work was the assessment of the relationship between FE traits (FCR and RFI) and feeding behaviour assessed by the number of feed demands over a given period and the proportion of the demands made before noon. The daily profile of feeding behaviour during the three FI measurement periods is presented in Figure 4A. Over the three periods, feeding behaviour changed over the hours (*P* < 0.001). Nocturnal demands were only occasional, as expected, because demands were rewarded only during the daylight hours. Even if limited re-rankings were observed between lines, the daily feed demand profiles remained similar between fi_1_ and fi_2,_ with, on average, 37-72% and 56-80% of the feed demands occurring in the morning. The morning feeding activity of fish is even more pronounced in fi_3,_ with 76-90% of demands recorded before noon (Figure 4A). There is limited information on how these seasonal or circannual rhythms influence feeding behaviour in fish, making it complex to give a straightforward conclusion about feeding activities and associated seasonal changes (Assan et al., 2021). Nevertheless, we can reasonably assume that the seasonal difference between the fi periods (autumn for fi1 and spring-summer for fi2 and fi3) may, at least partially, explain the changes observed in the daily feeding profile.

**Figure 4.**
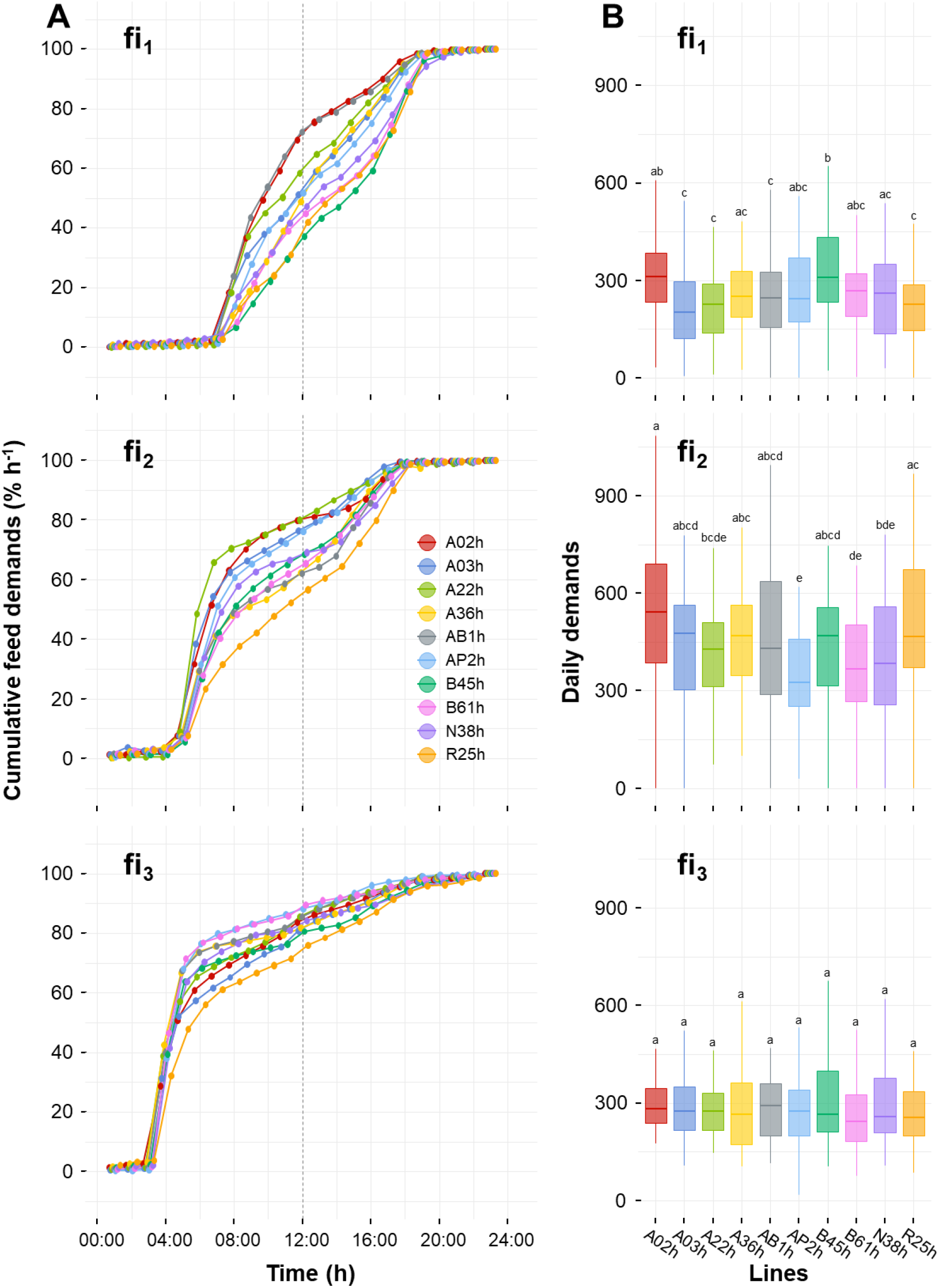
(A) Feeding behaviour expressed as the cumulative feed demands (% h^-1^) in the ten isogenic lines fed using self-feeders: from d 129 to d 161 (fi_1_), from d 301 to d 335 (fi_2_) and from d 388 to d 415 (fi_3_). In order to facilitate reading, values are the means of three replicates and SD were not represented. (B) Daily demands of the ten isogenic lines over fi_1_, fi_2_ and fi_3_ periods. Letters denote significant differences in daily feed demands between lines.

Significant differences in the daily profile of feed demands among the ten isogenic lines were found for the first FI measurement periods (fi_1_ and fi_2_; *P* ⩽ 0.001). Significant interactions between lines and time of the day were found (*P* ⩽ 0.001), suggesting that the effects of time of day were line-dependent. In fi_3_, daily feeding activity remained similar between lines (*P* = 0.421), while the number of feed demands per day remained constant between lines (*P* = 0.227, Figure 4B). Mambrini et al. (2004) highlighted, in brown trout, that a line selected for growth over five generations exhibited a more important morning feeding activity than the control line, maintained under the same rearing conditions, even if the variations in feeding activity between lines were not constant over time. Here, we did not find significant correlations between the proportion of the feed demand occurring in the morning and the TGC (Figure 3), suggesting that growth performances in isogenic lines are not related to the feeding profile of the fish.

In our study, the daily distribution of feed demands and their number were recorded as phenotypes for feeding behaviour. Indeed, the self-feeders allow continuous recording of feed demands (rewarded or not, as described in Section 2.4.2) without disturbing the behaviour of the fish group in the tank. Interestingly, we found significant positive correlations between RFI calculated from the first feed intake measurement period and feeding behaviour, assessed as the number of feed demands (*i*.*e*. rewarded, unrewarded and both of them) (r = 0.42-0.49, *P* ⩽ 0.022; Figure 3). It is interesting to note that the fact that the requests are rewarded or not has no influence on the correlations with the RFI. While the number of rewarded demands reflects FI, this is, by definition, not the case for unrewarded demands. Thus, we can hypothesise that this latter phenotype better reflects the motivation of fish for food (i.e. appetite). In other words, in the described rearing conditions, the most feed efficient isogenic lines (lower FCR and RFI) were the ones that requested less feed via the self-feeders and presumably had less appetite. In fi_3_, we found significant negative correlations between FCR or RFI and the proportion of the feed demands done in the morning (r = -0.41 and -0.38, *P* ⩽ 0.038; Figure 3), meaning that the most efficient fish had a higher feeding activity in the morning.

The assessment of feeding behaviour in fish is done through various criteria (e.g., number of feed demands, feeding latency, feeding consistency and total feeding time), making comparisons between studies difficult. Nevertheless, our results are in accordance with previous results in African catfish (Martins et al., 2006) and Nile tilapia (Martins et al., 2011a) that found significant positive phenotypic correlations between feeding latency, feeding time, and the number of feeding acts, and RFI while they did not find significant correlations with FCR. In another study in Nile tilapia, Martins et al. (2011b) found that the correlations between feeding behaviour, measured using the same traits as described above, and RFI depends on the age of the fish. Mas-Muñoz et al. (2011) also highlighted that consistency in feeding activity might be related to FE in common sole *Solea solea*. They found that fish which feed more consistently over time (within the day and over days) showed higher FI and growth but tend to be less feed efficient.

Overall, the feeding behaviour differences we observed between isogenic lines suggest scope for improvement in our knowledge of the genetic basis of fish feeding behaviour. There is growing evidence of a relationship between FE traits, especially RFI and feeding behaviour traits in fish, and this study confirms such relationships in rainbow trout while highlighting the need for repeated measurements over time. Therefore, special attention should be paid to behavioural traits that may be of interest in breeding programs for fish.

To conclude, we highlighted, using repeated measurements of FE traits (FCR and RFI), that the growth performances observed during FD periods are not easily linked to variations in FE measured in rainbow trout, suggesting that such indirect criteria for FE are not appropriate in selection. Interestingly, we highlighted some relationships between FE and feeding behaviour (*i*.*e*. number of feed demands over a period and the proportion of the demands made before noon) while the use of isogenic lines demonstrated the genetic basis of feeding behaviour. Overall, these results support the interest in exploring phenotypes of feeding behaviour as potential proxies of FE for improving RFI in rainbow trout through selective breeding.

## Acknowledgements

The authors thank the staff of the INRAE experimental facilities (PEIMA, INRAE, 2021, Fish Farming systems Experimental Facility, DOI: 10.15454/1.5572329612068406E12, Sizun, France).

## References

Ali, M., Nicieza, A., Wootton, R.J., 2003. Compensatory growth in fishes: a response to growth depression. Fish Fish. 4, 147–190. https://doi.org/10.1046/j.1467-2979.2003.00120.x

Assan, D., Huang, Y., Mustapha, U.F., Addah, M.N., Li, G., Chen, H., 2021. Fish Feed Intake, Feeding Behavior, and the Physiological Response of Apelin to Fasting and Refeeding. Front. Endocrinol. (Lausanne). 12, 1–12. https://doi.org/10.3389/fendo.2021.798903

Azzaydi, M., Rubio, V.C., López, F.J.M., Sánchez-Vázquez, F.J., Zamora, S., Madrid, J.A., 2007. Effect of restricted feeding schedule on seasonal shifting of daily demand-feeding pattern and food anticipatory activity in European sea bass (Dicentrarchus labrax L.). Chronobiol. Int. 24, 859–874. https://doi.org/10.1080/07420520701658399

Besson, M., Allal, F., Chatain, B., Vergnet, A., Clota, F., Vandeputte, M., 2019. Combining individual phenotypes of feed intake with genomic data to improve feed efficiency in sea bass. Front. Genet. 10, 1–14. https://doi.org/10.3389/fgene.2019.00219

Besson, M., Komen, H., Rose, G., Vandeputte, M., 2020. The genetic correlation between feed conversion ratio and growth rate affects the design of a breeding program for more sustainable fish production. Genet. Sel. Evol. 52, 5. https://doi.org/10.1186/s12711-020-0524-0

Boujard, T., Labbé, L., Aupérin, B., 2002. Feeding behaviour, energy expenditure and growth of rainbow trout in relation to stocking density and food accessibility. Aquac. Res. 33, 1233–1242. https://doi.org/10.1046/j.1365-2109.2002.00755.x

Bowyer, P.H., El-Haroun, E.R., Salim, H.S., Davies, S.J., 2020. Benefits of a commercial solid-state fermentation (SSF) product on growth performance, feed efficiency and gut morphology of juvenile Nile tilapia (Oreochromis niloticus) fed different UK lupin meal cultivars. Aquaculture 523. https://doi.org/10.1016/j.aquaculture.2020.735192

Cantalapiedra-Hijar, G., Faverdin, P., Friggens, N.C., Martin, P., 2020. Efficience Alimentaire : comment mieux la comprendre et en faire un élément de durabilité de l’élevage. INRAE Prod. Anim. 33, 235–248. https://doi.org/10.20870/productions-animales.2020.33.4.4594

Chavanne, H., Janssen, K., Hofherr, J., Contini, F., Haffray, P., Aquatrace Consortium, Komen, H., Nielsen, E.E., Bargelloni, L., 2016. A comprehensive survey on selective breeding programs and seed market in the European aquaculture fish industry. Aquac. Int. 24, 1287–1307. https://doi.org/10.1007/s10499-016-9985-0

Clarke, A., Johnston, N.M., 1999. Scaling of metabolic rate with body mass and temperature in teleost fish. J. Anim. Ecol. 68, 893–905. https://doi.org/https://doi.org/10.1046/j.1365-2656.1999.00337.x

da Silva, R.F., Kitagawa, A. Francisco, Sánchez Vázquez, F.J., 2016. Dietary self-selection in fish: a new approach to studying fish nutrition and feeding behavior. Rev. Fish Biol. Fish. 26, 39–51. https://doi.org/10.1007/s11160-015-9410-1

de Verdal, H., Komenc, H., Quillet, E., Chatain, B., Allal, F., Benzie, J.A.H., Vandeputte, M., 2018a. Improving feed efficiency in fish by selective breeding: a review. Rev. Aquac. 10, 833–851.

de Verdal, H., Mekkawy, W., Lind, C.E., Vandeputte, M., Chatain, B., Benzie, J.A.H., 2017. Measuring individual feed efficiency and its correlations with performance traits in Nile tilapia, Oreochromis niloticus. Aquaculture 468, 489–495. https://doi.org/10.1016/j.aquaculture.2016.11.015

de Verdal, H., O’Connell, C.M., Mekkawy, W., Vandeputte, M., Chatain, B., Bégout, M.L., Benzie, J.A.H., 2019. Agonistic behaviour and feed efficiency in juvenile Nile tilapia Oreochromis niloticus. Aquaculture 505, 271–279. https://doi.org/10.1016/j.aquaculture.2019.02.067

de Verdal, H., Vandeputte, M., Mekkawy, W., Chatain, B., Benzie, J.A.H., 2018b. Quantifying the genetic parameters of feed efficiency in juvenile Nile tilapia Oreochromis niloticus. BMC Genet. 19, 105. https://doi.org/10.1186/s12863-018-0691-y

Doupé, R.G., Lymbery, A.J., 2004. Indicators of genetic variation for feed conversion efficiency in black bream. Aquac. Res. 35, 1305–1309. https://doi.org/10.1111/j.1365-2109.2004.01128.x

Dupont-Prinet, A., Chatain, B., Grima, L., Vandeputte, M., Claireaux, G., McKenzie, D.J., 2010. Physiological mechanisms underlying a trade-off between growth rate and tolerance of feed deprivation in the European sea bass (Dicentrarchus labrax). J. Exp. Biol. 213, 1143–1152. https://doi.org/10.1242/jeb.037812

Dvergedal, H., Galloway, T., Sharma, S., Juarez, M., 2022. Verifying the relationship between d 13 C isotope profile variables and individual feed conversion ratio in large rainbow trout (Oncorhynchus mykiss). Aquaculture 558, 738355. https://doi.org/10.1016/j.aquaculture.2022.738355

El-Sayed, A.-F.M., 2002. Effects of stocking density and feeding levels on growth and feed efficiency of Nile tilapia (Oreochromis niloticus L.) fry. Aquac. Res. 33, 621–626. https://doi.org/10.1046/j.1365-2109.2002.00700.x

FAO, 2020. The State of World Fisheries and Aquaculture 2020, The State of World Fisheries and Aquaculture 2020. FAO, Roma, Italy. https://doi.org/10.4060/ca9229en

Franěk, R., Baloch, A.R., Kašpar, V., Saito, T., Fujimoto, T., Arai, K., Pšenička, M., 2020. Isogenic lines in fish: a critical review. Rev. Aquac. 12, 1412–1434. https://doi.org/10.1111/raq.12389

Gjedrem, T., 2010. The first family-based breeding program in aquaculture. Rev. Aquac. 2, 2– 15. https://doi.org/10.1111/j.1753-5131.2010.01011.x

Grima, L., 2010. Vers une amélioration de l’efficacité alimentaire chez le poisson. Agro Paris Tech.

Grima, L., Chatain, B., Ruelle, F., Vergnet, A., Launay, A., Mambrini, M., Vandeputte, M., 2010a. In search for indirect criteria to improve feed utilization efficiency in sea bass (Dicentrarchus labrax). Part II: Heritability of weight loss during feed deprivation and weight gain during re-feeding periods. Aquaculture 302, 169–174. https://doi.org/10.1016/j.aquaculture.2010.02.016

Grima, L., Quillet, E., Boujard, T., Robert-Granié, C., Chatain, B., Mambrini, M., 2008. Genetic variability in residual feed intake in rainbow trout clones and testing of indirect selection criteria. Genet. Sel. Evol. 40, 607–624. https://doi.org/10.1051/gse

Grima, L., Vandeputte, M., Ruelle, F., Vergnet, A., Mambrini, M., Chatain, B., 2010b. In search for indirect criteria to improve residual feed intake in sea bass (Dicentrarchus labrax). Part I: Phenotypic relationship between residual feed intake and body weight variations during feed deprivation and re-feeding periods. Aquaculture 300, 50–58. https://doi.org/10.1016/j.aquaculture.2010.01.003

Henryon, M., Jokumsen, A., Berg, P., Lund, I., Pedersen, P.B., Olesen, N.J., Slierendrecht, W.J., 2002. Genetic variation for growth rate, feed conversion efficiency, and disease resistance exists within a farmed population of rainbow trout (Aquaculture (2002) 209 (59-76) PII: S0044848601007293). Aquaculture 209, 59–76. https://doi.org/10.1016/S0044-8486(02)00607-5

Iwama, G.K., Tautz, A.F., 1981. A simple growth model for salmonids in hatcheries. Can. J. Fish. Aquat. Sci. 38, 649–656. https://doi.org/10.1139/f81-087

Jobling, M., 2002. Environmental Factors and Rates of Development and Growth, in: Hart, P.J.B., Reynolds, J.D. (Eds.), Handbook of Fish Biology and Fisheries, Wiley Online Books. pp. 97–122. https://doi.org/https://doi.org/10.1002/9780470693803.ch5

Jobling, M., Covès, D., Damsgard, B., Kristiansen, H.R., Koskela, J., Petursdottir, T.E., Kadri, S., Gudmundsson, O., 2001. Techniques for Measuring Feed Intake, in: Houlihan, D., Boujard, T., Jobling, M. (Eds.), Food Intake in Fish. Wiley-Blackwell, Oxford, pp. 49–87.

Just, P.N., Köllner, B., Slater, M.J., 2021. Video surveillance methods to evaluate individual feeding response in rainbow trout (Oncorhynchus mykiss, Walbaum)—implications for feeding regime optimisation. Aquac. Int. 29, 999–1013. https://doi.org/10.1007/s10499-021-00671-z

Kause, A., Kiessling, A., Martin, S.A.M., Houlihan, D., Ruohonen, K., 2016. Genetic improvement of feed conversion ratio via indirect selection against lipid deposition in farmed rainbow trout (Oncorhynchus mykiss Walbaum). Br. J. Nutr. 116, 1656–1665. https://doi.org/10.1017/S0007114516003603

Kause, A., Tobin, D., Houlihan, D.F., Martin, S.A.M., Mäntysaari, E.A., Ritola, O., Ruohonen, K., 2006. Feed efficiency of rainbow trout can be improved through selection: Different genetic potential on alternative diets. J. Anim. Sci. 84, 807–817. https://doi.org/10.2527/2006.844807x

Kelly, A.K., McGee, M., Crews, D.H., Fahey, A.G., Wylie, A.R., Kenny, D.A., 2010. Effect of divergence in residual feed intake on feeding behavior, blood metabolic variables, and body composition traits in growing beef heifers. J. Anim. Sci. 88, 109–123. https://doi.org/10.2527/jas.2009-2196

Knap, P.W., Kause, A., 2018. Phenotyping for genetic improvement of feed efficiency in fish: Lessons from pig breeding. Front. Genet. 9, 184. https://doi.org/10.3389/fgene.2018.00184

Koch, R.M., Swiger, L.A., Chambers, D., Gregory, K.E., 1963. Efficiency of feed use in beef cattle. J. Anim. Sci. 22, 486–494. https://doi.org/10.2527/jas1963.222486x

Komen, H., Thorgaard, G.H., 2007. Androgenesis, gynogenesis and the production of clones in fishes: A review. Aquaculture 269, 150–173. https://doi.org/10.1016/j.aquaculture.2007.05.009

Lallias, D., Quillet, E., Bégout, M.-L., Aupérin, B., Khaw, H.L., Millot, S., Valotaire, C., Kernéis, T., Labbé, L., Prunet, P., Dupont-Nivet, M., 2017. Genetic variability of environmental sensitivity revealed by phenotypic variation in body weight and (its) correlations to physiological and behavioral traits. PLoS One 12, e0189943. https://doi.org/10.1371/journal.pone.0189943

Li, M.H., Robinson, E.H., Bosworth, B.G., 2005. Effects of periodic feed deprivation on growth, feed efficiency, processing yield, and body composition of channel catfish Ictalurus punctatus. J. World Aquac. Soc. 36, 444–453. https://doi.org/10.1111/j.1749-7345.2005.tb00392.x

Mambrini, M., Sanchez, M.P., Chevassus, B., Labbé, L., Quillet, E., Boujard, T., 2004. Selection for growth increases feed intake and affects feeding behavior of brown trout. Livest. Prod. Sci. 88, 85–98. https://doi.org/10.1016/j.livprodsci.2003.10.005

Martins, C.I.M., Conceição, L.E.C., Schrama, J.W., 2011a. Feeding behavior and stress response explain individual differences in feed efficiency in juveniles of Nile tilapia Oreochromis niloticus. Aquaculture 312, 192–197. https://doi.org/10.1016/j.aquaculture.2010.12.035

Martins, C.I.M., Conceição, L.E.C., Schrama, J.W., 2011b. Consistency of individual variation in feeding behaviour and its relationship with performance traits in Nile tilapia Oreochromis niloticus. Appl. Anim. Behav. Sci. 133, 109–116. https://doi.org/10.1016/j.applanim.2011.05.001

Martins, C.I.M., Trenovski, M., Schrama, J.W., Verreth, J.A.J., 2006. Comparison of feed intake behaviour and stress response in isolated and non-isolated African catfish. J. Fish Biol. 69, 629–636. https://doi.org/10.1111/j.1095-8649.2006.01121.x

Mas-Muñoz, J., Komen, H., Schneider, O., Visch, S.W., Schrama, J.W., 2011. Feeding behaviour, swimming activity and boldness explain variation in feed intake and growth of sole (Solea solea) reared in captivity. PLoS One 6, 1–9. https://doi.org/10.1371/journal.pone.0021393

Mattila, J., Koskela, J., Pirhonen, J., 2009. The effect of the length of repeated feed deprivation between single meals on compensatory growth of pikeperch Sander lucioperca. Aquaculture 296, 65–70. https://doi.org/10.1016/j.aquaculture.2009.07.024

Mente, E., Pierce, G.J., Santos, M.B., Neofitou, C., 2006. Effect of feed and feeding in the culture of salmonids on the marine aquatic environment: A synthesis for European aquaculture. Aquac. Int. 14, 499–522. https://doi.org/10.1007/s10499-006-9051-4

Millot, S., Bégout, M.L., 2009. Individual fish rhythm directs group feeding: A case study with sea bass juveniles (Dicentrarchus labrax) under self-demand feeding conditions. Aquat. Living Resour. 22, 363–370. https://doi.org/10.1051/alr/2009048

Millot, S., Péan, S., Labbé, L., Kerneis, T., Quillet, E., Dupont-Nivet, M., Bégout, M.-L., 2014. Assessment of genetic variability of fish personality traits using rainbow trout isogenic lines. Behav. Genet. 44, 383–393. https://doi.org/10.1007/s10519-014-9652-z

Montanholi, Y.R., Swanson, K.C., Palme, R., Schenkel, F.S., McBride, B.W., Lu, D., Miller, S.P., 2010. Assessing feed efficiency in beef steers through feeding behavior, infrared thermography and glucocorticoids. Animal 4, 692–701. https://doi.org/10.1017/S1751731109991522

Nikki, J., Pirhonen, J., Jobling, M., Karjalainen, J., 2004. Compensatory growth in juvenile rainbow trout, Oncorhynchus mykiss (Walbaum), held individually. Aquaculture 235, 285–296. https://doi.org/10.1016/j.aquaculture.2003.10.017

Oh, S.Y., Noh, C.H., Cho, S.H., 2007. Effect of restricted feeding regimes on compensatory growth and body composition of red sea bream, Pagrus major. J. World Aquac. Soc. 38, 443–449. https://doi.org/10.1111/j.1749-7345.2007.00116.x

Quillet, E., Dorson, M., Le Guillou, S., Benmansour, A., Boudinot, P., 2007a. Wide range of susceptibility to rhabdoviruses in homozygous clones of rainbow trout. Fish Shellfish Immunol. 22, 510–519. https://doi.org/10.1016/j.fsi.2006.07.002

Quillet, E., Le Guillou, S., Aubin, J., Labbé, L., Fauconneau, B., Médale, F., 2007b. Response of a lean muscle and a fat muscle rainbow trout (Oncorhynchus mykiss) line on growth, nutrient utilization, body composition and carcass traits when fed two different diets. Aquaculture 269, 220–231. https://doi.org/10.1016/j.aquaculture.2007.02.047

R Development Core Team, 2020. R: A language and environment for statistical computing.

Read, P., Fernandes, T., 2003. Management of environmental impacts of marine aquaculture in Europe, in: Aquaculture. Elsevier, pp. 139–163. https://doi.org/10.1016/S0044-8486(03)00474-5

Silverstein, J.T., 2006. Relationships among Feed Intake, Feed Efficiency, and Growth in Juvenile Rainbow Trout. N. Am. J. Aquac. 68, 168–175. https://doi.org/10.1577/a05-010.1

Talbot, C., Higgins, P.J., 1983. A radiographic method for feeding studies on fish using metallic iron powder as a marker. J. Fish Biol. 23, 211–220. https://doi.org/10.1111/j.1095-8649.1983.tb02896.x

Thodesen, J., Grisdale-helland, B., Helland, J., Gjerde, B., 1999. Feed intake, growth and feed utilization of offspring from wild and selected Atlantic salmon ž Salmo salar /. Aquaculture 180, 237–246.

Troell, M., Naylor, R.L., Metian, M., Beveridge, M., Tyedmers, P.H., Folke, C., Arrow, K.J., Barrett, S., Crépin, A.S., Ehrlich, P.R., Gren, Å., Kautsky, N., Levin, S.A., Nyborg, K., Österblom, H., Polasky, S., Scheffer, M., Walker, B.H., Xepapadeas, T., De Zeeuw, A., 2014. Does aquaculture add resilience to the global food system? Proc. Natl. Acad. Sci. U. S. A. https://doi.org/10.1073/pnas.1404067111

Weatherley, A.H., Gill, H.S., 1981. Recovery growth following periods of restricted rations and starvation in rainbow trout Salmo gairdneri Richardson. J. Fish Biol. 18, 195–208. https://doi.org/10.1111/j.1095-8649.1981.tb02814.x

Wu, L., Xie, S., Zhu, X., Cui, Y., Wootton, R.J., 2002. Feeding dynamics in fish experiencing cycles of feed deprivation: A comparison of four species. Aquac. Res. 33, 481–489. https://doi.org/10.1046/j.1365-2109.2002.00733.x

Xu, X.L., Fontaine, P., Mélard, C., Kestemont, P., 2001. Effects of dietary fat levels on growth, feed efficiency and biochemical compositions of Eurasian perch Perca fluviatilis. Aquac. Int. 9, 437–449. https://doi.org/10.1023/A:1020597415669

Yilmaz, Y., Arabaci, M., 2010. The influence of stocking density on growth and feed efficiency in gilthead seabream, Sparus aurata. J. Anim. Vet. Adv. 9, 1280–1284.

Zhou, C., Xu, D., Lin, K., Sun, C., Yang, X., 2018. Intelligent feeding control methods in aquaculture with an emphasis on fish: a review. Rev. Aquac. 10, 975–993. https://doi.org/10.1111/raq.12218

